# PRECISE EXOME ANALYSIS OF BLASTOCYST BIOPSY SCALE SAMPLES USING PRIMARY TEMPLATE-DIRECTED AMPLIFICATION

**DOI:** 10.1101/2024.10.29.620888

**Authors:** Alina Samitova, Vera Belova, Iuliia Vasiliadis, Zhanna Repinskaia, Tatiana Gorodnicheva, Evgeny Romanov, Mariam Pogosyan, Emil Gaysin, Tatyana Nazarenko, Denis Rebrikov, Dmitriy Korostin

## Abstract

This study evaluates primary template-directed amplification (PTA) for whole exome sequencing (WES) on small fibroblast cell groups, mimicking the limited cell quantities typical of trophectoderm embryo biopsies. PTA’s consistent amplification reduces allelic dropout (ADO) and impoves uniform coverage, overcoming challenges associated with conventional methods such as multiple displacement amplification (MDA). Using fibroblast samples alongside well-characterised genomic references (E701, NA12878), we benchmarked PTA-WES, achieving 97.5% target region coverage at 10x, meeting American College of Medical Genetics and Genomics (ACMG) standards. Preliminary results from embryo biopsies sequenced with PTA-WES showed a median coverage of 102x, significantly improving upon the variability and coverage gaps observed in MDA-WES. The findings support PTA’s potential to enhance the clinical applicability of WES for preimplantation genetic testing for monogenic disorders (PGT-M), expanding capabilities to detect inherited and de novo mutations in embryos. Further optimisation and variant detection analyses are planned to evaluate PTA’s robustness for routine clinical use.

## Introduction

Preimplantation genetic testing (PGT) is a crucial tool in assisted reproductive technologies, allowing for the selection of embryos free from genetic abnormalities before implantation. Previously, many methods were unavailable for the analysis of trophectoderm biopsies from embryos. However, with improvements in whole genome amplification (WGA) protocols, it has become possible to obtain sufficient starting material from as little as 6–7 picograms of DNA from a single cell, overcoming prior limitations of existing methods and expanding the potential for embryo genome analysis (1-5). Among current techniques, multiple displacement amplification (MDA) and Multiple annealing and looping-based amplification cycles (MALBAC) have shown significant promise, particularly in PGT-A (testing for aneuploidies), due to its ability to amplify large quantities of DNA from minimum samples (6-8). Nevertheless whole exome sequencing (WES), a powerful tool for detecting a wide range of genetic variants associated with inherited disorders, remains largely unavailable in PGT. Despite its advantages, MDA’s uneven amplification of DNA leads to coverage gaps, limiting the effectiveness of WES for preimplantation genetic testing for monogenic disorders (PGT-M) (9-12).

Currently, PGT-M involves the collection of maternal, paternal, and control samples, which are analyzed alongside amplified genomic DNA from trophectoderm biopsies to determine the mutation status of each embryo, using methods such as PCR with NGS or Sanger sequencing, karyomapping, haplarithmisis, etc. (12-14). Several studies highlight the critical role of PGT-M in the genetic analysis of embryos. For example, one study specifically highlighted the use of PGT-M to prevent transmission of genetic conditions such as Marfan syndrome through preimplantation genetic testing, where a PGT-M protocol was developed and tested using multiplex fluorescent PCR and mini-sequencing (15). There is also a known case of PGT for Meckel syndrome, where the WGA products of each embryo were subjected to Sanger sequencing for direct identification of variant sites, and haplotyping analysis was conducted using SNP markers (16). Although these approaches are effective for detecting target variants, they do not provide the same resolution as whole exome sequencing and do not allow for reliable identification of *de novo* variants. Some studies have provided data on WGS and WES of single cells (17,18). In addition, one study highlighted that exome sequencing can reveal clinically significant information about preimplantation embryos that may not be detectable in parental genomes (19). However, limitations in current WGA techniques such as allele drop-out (ADO), locus drop-out (LDO), chimeric DNA molecules, base replication errors and unevenness in amplification often prevent these techniques from achieving the high coverage required by standard of the American College of Medical Genetics and Genomics (ACMG) for accurate variant detection (20). It is also necessary to conduct not only mutation site detection but also SNP linkage analysis to ensure accuracy. Therefore improved ability to sequence high-quality exomes directly from trophectoderm biopsies could significantly increase the speed and accessibility of PGT-M, providing a broader range of genetic insights.

A potential solution to this challenge is the introduction of primary template-directed amplification (PTA), a novel method that offers more uniform WGA from small amounts of material, including preimplantation biopsy samples (21-23). PTA more evenly amplifies both alleles in the same cell, resulting in significantly diminished allelic dropout (ADO) and skewing. To evaluate the effectiveness of PTA for exome analysis, we conducted a study using fibroblast cultures with a well-characterized in-lab reference genome E701 and Platinum Genome DNA sample NA12878 (24, 25). This research aims to assess the applicability of PTA for expanding the capabilities of PGT, making WES a feasible and reliable option in clinical settings.

## Materials and Methods

### Sample collection

Human fibroblasts from a reference sample E701 from our laboratory were used for this study. They were carefully thawed from cryostorage and cultured in Dulbecco’s modified Eagle’s medium (DMEM) supplemented with 10% fetal bovine serum (FBS), penicillin-streptomycin (50 U/ml) and L-glutamine (2mM) to promote cell growth and maintain optimal conditions for cell proliferation. Once cell number was reached, fibroblasts were detached from the culture flask surface using 0.25% trypsin-EDTA solution according to a standard trypsinisation protocol to ensure cell viability and maintain consistent cell morphology for downstream applications.The fibroblasts were divided into groups of 4, 10, 16, and 25 cells, which were then placed in 0.2 ml tubes. Each group was prepared in triplicate. In addition, reference genomic DNA NA12878 and our in-lab reference DNA sample E701 was used at concentrations equivalent to 10 and 40 genomes.

This study also presents WES results from trophectoderm embryo biopsies. All embryo samples were donated for research purposes and provided by the V.I. Kulakov National Medical Research Center for Obstetrics, Gynecology, and Perinatology under the category of not suitable for implantation. The embryos were created by via intracytoplasmic sperm injection (ICSI). Embryos on days 5-6 were biopsied according to standard operating procedures (SOP) in Kulakov Center. Each biopsy was collected in a 200 µL PCR tube containing 3 µL of cell buffer.

### WGA

PTA was performed according to the manufacturer’s instructions (BioSkryb Genomics). After PTA, DNA was purified using 2X Kapa Pure Beads (Roche). The DNA yield was quantified with the Qubit dsDNA HS Assay system (Life Technologies) and its quality was assessed by 1.5% agarose gel electrophoresis. MDA was performed using QIAGEN REPLI-g kits according to the manufacturer’s instructions (Qiagen).

### Sample Preparation and Exome Sequencing

For each library, 500 ng of PTA product or genomic DNA (for NA12878 and E701) was sheared using a Covaris LE220 according to the manufacturer’s protocol, followed by size selection with Kapa Pure Beads (Roche) to achieve a fragment distribution peak at ∼250 bp. DNA libraries were prepared using the MGI Universal DNA Library Prep Set, followed by final amplification with 8 PCR cycles. The concentration of the prepared libraries was measured using the Qubit Flex system (ThermoFisher) and the dsDNA HS Assay Kit. Quality control of the DNA libraries was performed using high-sensitivity analysis on the Bioanalyzer 2100 system (Agilent Technologies). Then, we pooled 900 ng of each of the 16 libraries into a single pool which was concentrated using a SpeedVac concentrator (ThermoFisher) at 60°C. Exome enrichment of the pool with the Agilent SureSelect Human All Exon V8 probes was performed according to the RSMU_exome protocol (4). The pool was then circularized, downloaded into 4 flow cell lanes and sequenced paired-end mode on the DNBSEQ-G400 platform, using the DNBSEQ-G400RS High-throughput Sequencing Set PE100 kit, following the manufacturer’s instructions (MGI Tech).

### Genomic Data Analyses

The quality of the obtained fastq files was analysed using FastQC v0.11.9 (26). Based on the quality metrics, the fastq files were trimmed using BBDuk by BBMap v38.96 (27). Reads were aligned to the indexed reference genome GRCh38.p14 using bwa-mem2 v2.2.1 (28). SAM files were converted into BAM files and sorted using SAMtools v1.9 to check the percentage of the aligned reads (29). Based on the obtained BAM files, the quality metrics of exome enrichment and sequencing were calculated using Picard v2.22.4, and the number of duplicates was calculated using Picard MarkDuplicates v2.22.4 (30).

## Results

### 1. PTA-WGA of Genomic DNA and DNA from Fibroblast Cells

To assess the quality of material obtained after WGA using the PTA method, we utilized both genomic DNA and fibroblast cells. Fibroblast cells were grouped by cell count (4, 10, 16, and 25) to model the typical cell quantities obtained from blastocyst trophectoderm biopsies. PTA-WGA was successful in all tested samples with no observed outliers. Based on electrophoresis and concentration measurements all samples showed similar characteristics. On average, we obtained ∼1600 ng of DNA per sample after PTA-WGA and further purification using 2x Kapa Pure Beads (Roche).

### 2. Raw sequencing data and Pooling balance

We obtained between 190 and 249 million (M) reads per each of 16 samples (Figure 1). As shown in the graph, the amount of data for samples with a low number of cells was comparable to that of samples from other groups, indicating a well-balanced pooling. Pre-capture pooling can be challenging when using different sample types (e.g., genomic DNA or FFPE DNA), as variations in initial DNA quality can lead to inconsistent data yields post-enrichment. Therefore, this initial test of pooling different cell groups within a single pool was successful in achieving comparable data output across sample

**Figure 1.**
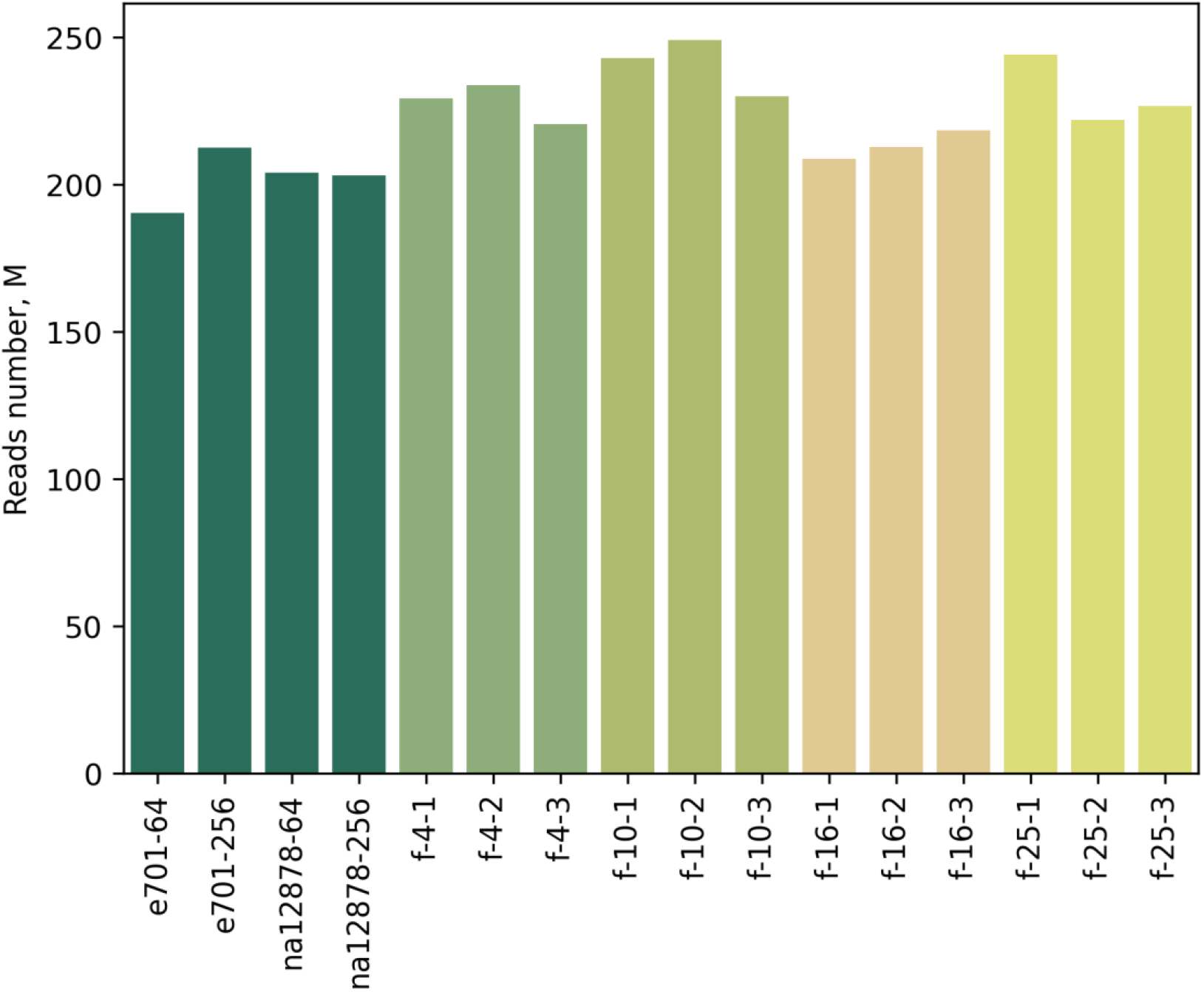
The barplot shows the number of reads per sample

### 3. Exome enrichment quality after PTA-WGA for fibroblast cells and genomic DNA

To assess exome sequencing quality after PTA-WGA, we grouped the samples as follows: samples originally taken with 4, 10, 16 and 25 fibroblast cells, and genomic DNA (E701 and NA12878). Coverage statistics were calculated using Picard and metrics were averaged across the sample groups. The results of Picard key metrics for all samples are presented in the Supplementary table 1. The percentage of on-target, off-target reads and duplicates for each sample group was calculated (Figure 2A). The distribution of these metrics did not reveal any particular dependence between groups of samples. However, it should be noted that in practical conditions it is not always possible to obtain ∼200 M reads per sample. We therefore downsampled each sample to 100 M reads. This process allows us to simulate conditions closer to basic exome data yield for clinical purposes. In this way, we can better estimate sequencing metrics for samples with less data. The graph (Figure 2B) shows that using 100 M reads per sample increased the average on-target percentage by 0.48% compared to the data shown in the graph (Figure 2 A) for samples with 200 million reads. This increase may be due to the reduction in duplicates observed with fewer reads.

**Figure 2.**
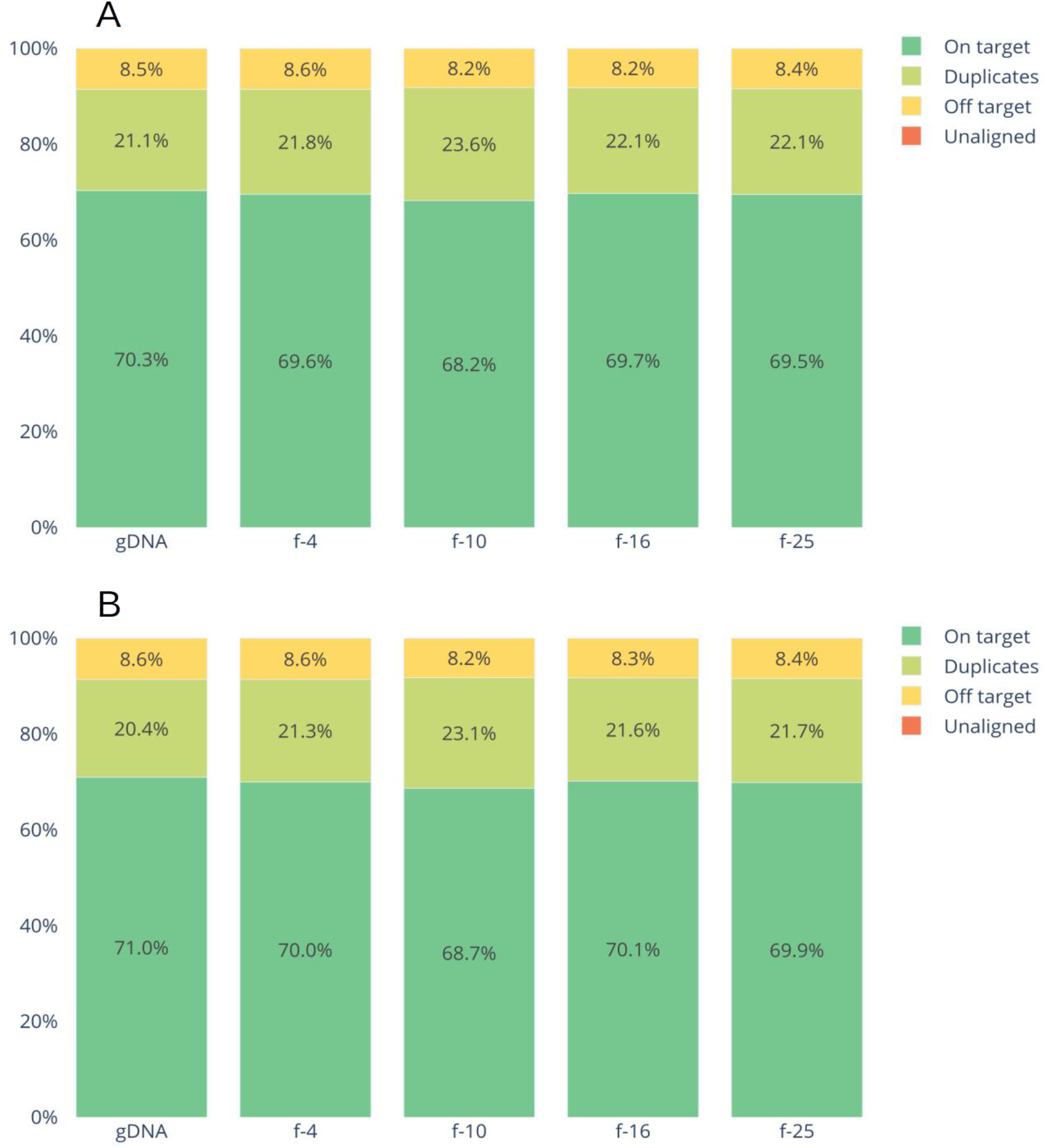
Barplots show the average on-targets, duplicates, and off-targets reads in each group for: (**A**) raw data, (**B**) downsampled data. The gDNA group included the laboratory standard sample e701 and the NA12878 reference sample, both of which were subjected to PTA in amounts of 64 pg and 256 pg, corresponding to approximately 10 and 40 genome equivalents.The f-4 group consisted of three replicate fibroblast samples, each containing four cells used for PTA. Similarly, the f-10 group consisted of three replicate fibroblast samples of ten cells each, while the f-16 group consisted of three replicate samples of sixteen cells each. Finally, the f-25 group consisted of three replicate fibroblast samples of twenty-five cells each for WGA.

The mean target coverage for all samples was 247x, with a median target coverage of 226x (Figure 3).

**Figure 3.**
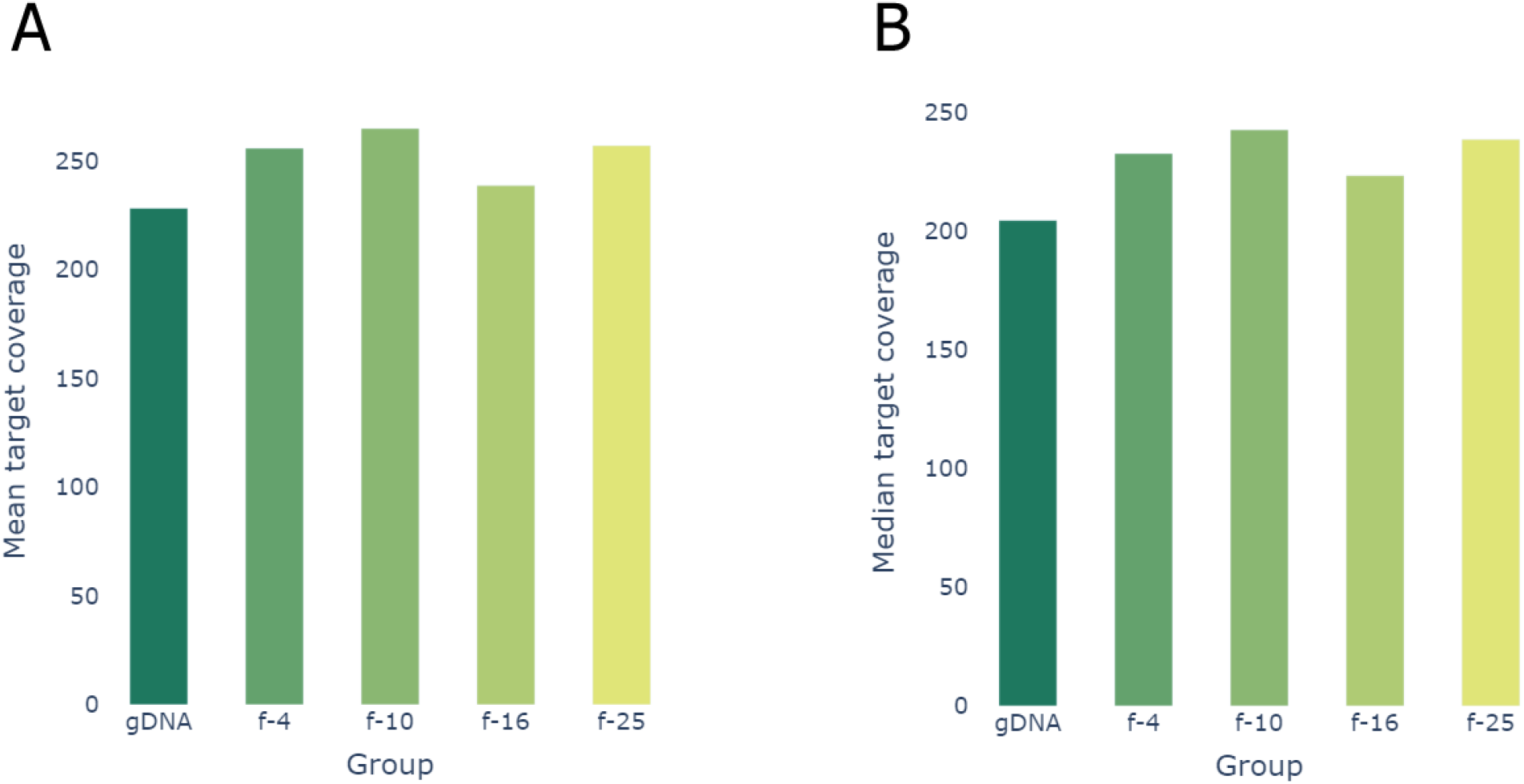
A) The average of the mean target coverage and B) the median target coverage of the samples that were analyzed without downsampling

Analysis of these coverage metrics showed no observable link between sequencing results and the type of starting material used. Furthermore, no dependency was noted between groups stratified by the number of cells used for PTA, indicating that variations in cell number in this experiment did not affect the consistency of coverage across samples.

The mean breadth, defined as the percentage of target regions covered at least x times per sample, for all samples in the pool was 97.5% (±0.26%SD) at 10x coverage depth, 96.25% (±0.59%SD) at 20x and 94.84% (±0.90%SD) at 30x. These parameters are characterized by a low standard deviation, indicating high homogeneity of the data (Figure 4).

**Figure 4.**
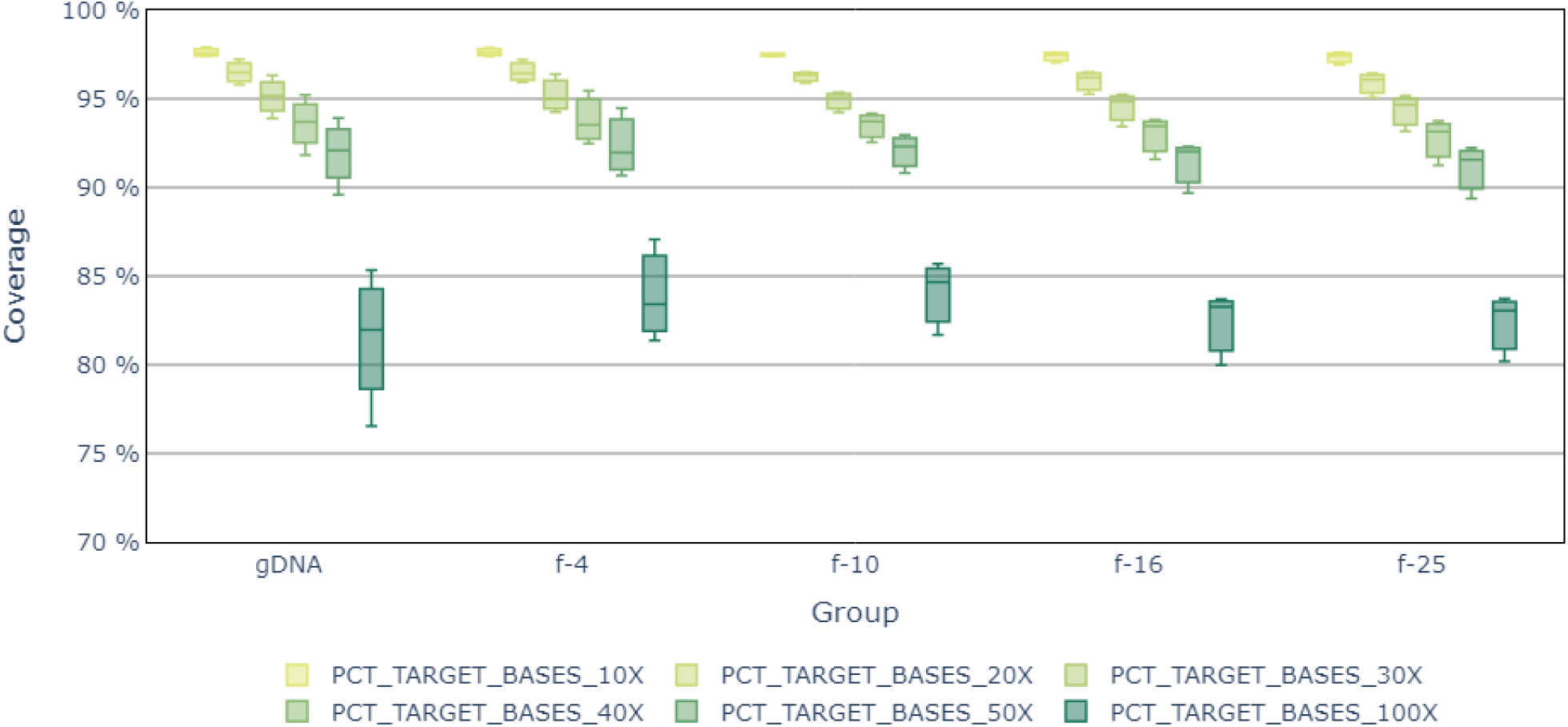
The average percentage of regions with 10x, 20x, 30x, 40x, 50x and 100x coverage depths for different groups of samples.

In this study, we observed an average of 1.28% (±0.03%SD) regions with zero coverage, which is fully consistent with the exome data typically obtained for samples isolated from blood (31).

### 4. ACMG Gene Coverage Analysis

We calculated the depth of coverage for 81 genes recommended by ACMG for the identification of pathogenic variants in clinical reporting (ACMG SF v3.2) (32). The average percentage of target regions covered at least 10x was over 98.4%. Further examination of the coverage distribution confirmed that the vast majority of target regions achieved uniform coverage in all samples (Supplementary table 2). Of the 81 clinically important genes analysed, 56 were fully covered at 100% across all samples. A further 11 genes achieved average coverage of 95-99% in all samples. However, coverage issues were observed for 6 genes, with an average coverage of around 80%. (Figure 5).

**Figure 5.**
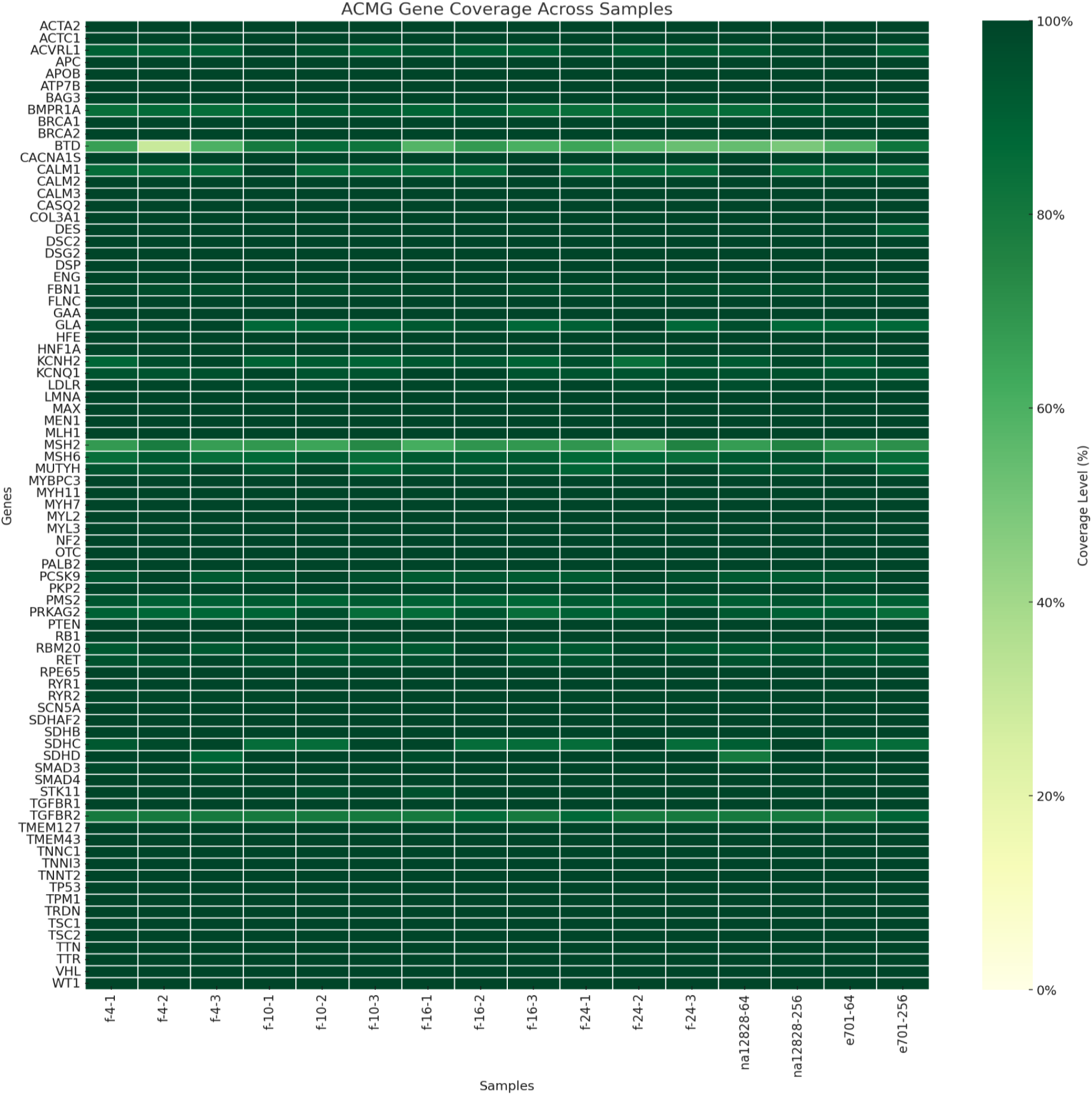
Variability in breadth of coverage for the American College of Medical Genetics and Genomics (ACMG) 81 genes among samples. Each row represents a gene, while each column corresponds to an individual sample.

### 5. PTA-WES vs MDA-WES for blastocyst trophectoderm embryo biopsies

Here we also present our preliminary results of an experiment using trophectoderm biopsies from blastocyst trophectoderm subjected to the PTA-WGA method. Seven different exomes from embryo biopsies samples were sequenced, resulting in 70-120 M reads per sample with mean coverage of 136x and median coverage of 102x with 85.71% of on-target reads. Coverage statistics across samples indicated that the percentage of target regions covered 10x ranged from 86.39% to 96.58%, with an average of 92.29% (±4.25%SD); for coverage at 20x it ranged from 77.73% to 94.75%, with an average of 86.93% (SD = 6.86%). The proportion of target regions with zero coverage remained low (1.27-2.23%, SD = 0.33%).

Additionally, in our previous experiments with six WES from blastocyst trophectoderm biopsies using WGA by MDA we encountered the problem of a high proportion of uncovered regions and a greater coverage unevenness. With the similar range of data yield per exome (70-160 M reads) the mean target coverage was 124x (±40.56%SD) but the median was only 6x (±9.97%SD) suggesting that many regions may not consistently reach adequate coverage in all samples. Percentage of regions covered 10x did not reach 95% and ranged only from 20.79% to 63.30% (SD = 16%) and the percentage of regions with zero coverage ranged from 11.78% to 55.13% (SD = 18.5%) demonstrating MDA-WES’s inability to achieve consistent target coverage.

## Discussion

In this study, we evaluated the quality of WES on small fibroblasts cell groups that underwent PTA-based WGA simulating the scale of trophectoderm embryo biopsies. To ensure a robust comparison, we included in-lab reference genomic DNA E701 and DNA of NA12878 Platinum genome sample.

The results demonstrate that PTA-based amplification generates sufficient amplified DNA material for library preparation and further exome enrichment regardless of the initial amount of fibroblast cells. After PTA step amplified products display a wide range (of ∼250 to >1,500 bp) of fragment sizes, as confirmed by gel electrophoresis results, suggesting that library preparation could bypass the fragmentation step. However, we chose to retain this step in the current study to avoid any potential loss of unique fragments. Fibroblasts DNA library samples containing varying initial cell counts (4, 10, 16, 25 cells) in this study were pre-capture multiplexed without compromising final data output. Low cell-count samples produced data yields similar to those of larger groups highlighting that PTA-WGA might support high-throughput sequencing in diverse cell input conditions. However we want to mention that in our previous experiments with trophectoderm embryo biopsies samples (data not shown), we encountered higher variability (5x times) in the amount of data obtained between samples within the same pool, despite identical preparation conditions. Probably for the embryo biopsy samples there are other factors that may influence pre-capture pooling balance, such as the quality of the embryo itself, its DNA integrity. Further research into embryo-specific characteristics that impact this stage of exome enrichment is needed.

Sequencing analysis results demonstrate that PTA enables consistent and efficient amplification across fibroblast single-cell genomes. Samples amplified by PTA achieved a mean percentage of target regions of 97.5% at 10x depth meeting the ACMG recommendations of 95% at 10x depth. The mean and median target coverage values were comparable to GIAB and E701 DNA exomes after PTA. Additionally, we observed that PTA-amplified samples contained only about 1.28% uncovered regions, aligning with the expected coverage gaps often encountered in exome sequencing due to low mappability genomic regions (31,33).

The primary quality metrics of PTA-amplified fibroblast samples at embryo biopsy scale meet established ACMG standards, showing its potential to clinical genetic analysis. Therefore we present coverage breadth results for the 81 genes recommended by the ACMG for pathogenic variant identification. The average coverage for over 67 genes exceeded 95%, demonstrating the efficacy of the PTA amplification method. These results indicate that PTA-based WGA meets initial requirements for further accurate variant detection for screening for monogenic disorders.

Our study is the first to achieve successful WES from trophectoderm human embryo biopsies. The comparative analysis of PTA-WES and MDA-WES on trophectoderm biopsies highlights significant advancements and limitations in PGT-M. The PTA-WGA approach for embryos demonstrated superior performance providing more uniform amplification, thus, higher median exome coverage (102x), low zero-coverage regions (mean 1.51%), and stable target region coverage at 10x and 20x depths (mean 92.29% and 86.93%, respectively). At this stage, we hypothesize that increasing the read output per sample could further improve coverage for WES in embryo biopsies, as several samples already exceeded 95% of target regions at 10x depth. These findings underscore the potential of PTA-based WGA to support robust WES in PGT-M, facilitating the detection of genetic variants. Our future work will include variant calling analyses to enhance our evaluation of PTA-WES. In contrast, the MDA-WGA method produced less consistent results, with substantial coverage gaps and a higher percentage of zero-coverage regions (up to 55.13%), underscoring the challenges associated with its use in PGT-M.

As per our study PTA by BioSkryb Genomics is a promising WGA tool for exome sequencing in PGT-M. PTA’s improved performance could expand the utility of WES in clinical applications by allowing for the detection of both inherited and de novo variants. Detecting de novo pathogenic variants is particularly valuable in PGT-M, as these variants may not be present in parental genomes and may be associated with severe developmental disorders. Despite these advancements, there are still challenges in integrating WES with PTA-WGA into standard PGT-M workflows. Variability in starting material quality and sample preparation can affect data output. Future research should continue to evaluate PTA’s long-term reliability in clinical applications and explore further optimization in both wet lab and computational methods.

## Supporting information

Supplementary table 1. Summary of key metrics for exome sequencing

Supplementary table 2. ACMG Gene Coverage Metrics

Supplementary table 3. Exome sequencing metrics for embryo biopsies undergoing WGA using PTA and MDA.

## Author Contributions

AS: designed, performed research and analyzed data; writing - original draft; writing - review & editing; VB: designed, performed research and analyzed data; supervision; writing - review & editing; YV: software and visualization; ZR: software and visualization; TG: methodology; provided experimental samples; ER: methodology; provided experimental samples; MP: provided experimental samples; EG: provided experimental samples; TA: provided experimental samples; DK: designed research; resources and funding acquisition; conceptualization; project administration; DR: funding acquisition;

## Ethics declarations

The appropriate institutional review board approval for this study was obtained from the Ethics Committee at the Pirogov Russian National Research Medical University.

